# Antiparallel Cell Circulation Emerging from Self-Aligned Tension Gradients

**DOI:** 10.1101/2025.07.28.667323

**Authors:** Ryunosuke Karimata, Hidenori Hashimura, Shuhei A. Horiguchi, Taihei Fujimori, Satoshi Sawai, Satoru Okuda

## Abstract

Active tissues exhibit diverse collective dynamics, yet the cell–cell interactions that generate ordered microscopic flows remain poorly understood. Here, we show that antiparallel cell circulation can emerge from self-aligned, polarity-dependent tension gradients. Using a minimal vertex model of confluent tissues, we studied polar cells that align their polarity with their own velocity and impose polarity-dependent tension gradients along cell–cell contacts, without relying on substrate traction. This behavior can be generalized as a minimal interaction in which forces transmitted between cells act with opposite signs, reminiscent of action–reaction forces, organizing cells into stable interlocking antiparallel lanes. In mixtures of motile and nonmotile cells, this circulation drives phase separation, in which motile cells spontaneously form persistent domains. Accordingly, we identified similar antiparallel circulation patterns in two-dimensional aggregates of *Dictyostelium discoideum*, supporting the biological relevance of the mechanism. Together, these results demonstrate that self-aligned tension gradients provide a robust and underappreciated route to dynamic microscopic pattern formation in multicellular systems.

## I. INTRODUCTION

Ordered patterns are a hallmark of active matter, appearing in diverse systems such as self-propelled colloids [1–3], active droplets [4,5], bacterial suspensions [6–9], and living cell assemblies [10–12]. Such systems can exhibit diverse spatiotemporal order [7,8,13,14], demonstrating how internally generated active forces can coordinate matter far from equilibrium. A natural mechanistic question is how these forces are exchanged and transmitted between constituents. This issue is especially relevant in densely packed biological assemblies, where the dominant mechanical interactions often arise not from interactions with the surrounding environment, but from force exchange and transmission via direct physical contacts between constituents. A simplified setting that highlights such contact-mediated force transmission is provided by motor-filament assays such as actin–myosin and microtubule–kinesin systems: cytoskeletal filaments linked by motor proteins slide past one another [15]. Similar contact-mediated force exchange also occurs at cell–cell junctions in confluent tissues. Yet how inter-constituent force exchange shapes spatiotemporal organization at the single-cell scale in confluent tissues remains poorly understood.

Epithelial tissues are a good example of cell-based active systems, exhibiting a variety of dynamic patterns, such as flocking [16–19] and vortex formation [19–23]. Often, cells migrate on frictional substrates or basement membranes, generating motion via multiple types of active driving forces. The emergence of patterns driven by self-propulsion at the cell–substrate interface has been widely investigated in both theoretical and experimental studies [22,24–28]. A distinct, less explored possibility arises from active interfacial tensions at cell–cell contacts, such as epithelial junctions [29–31]. Because interfacial tensions exert equal and opposite forces on the two neighboring cells, they obey action–reaction symmetry and provide a natural route to internally coordinated motion without requiring net external forces. Importantly, these tensions can become directionally biased by intracellular polarity, producing polarity-aligned interfacial tension gradients that drive directional cell migration and complex collective dynamics in confluent tissues [29–31].

Polarity regulation plays a central role in determining how mechanical interactions are translated into collective cell behavior. Two primary mechanisms have been considered: mutual alignment, in which neighboring cells directly coordinate their polarities [17,18,32,33], and self-alignment, where polarity aligns with each cell’s own movement [24,34–37]. Mutual alignment can directly prescribe the local arrangement of cell polarities at the single-cell level, whereas in self-alignment, polarity is generated intrinsically and does not by itself specify how neighboring cells arrange or orient [35,37,38]. In systems dominated by self-alignment, cell-scale order is therefore likely to emerge from additional intercellular mechanical interactions. Interfacial tension gradients offer a natural mechanism for such coupling: by biasing active tensions along cell–cell contacts, they generate directional yet force-balanced interactions that feed back onto cell rearrangements and motion. Whether these self-aligned tension gradients alone can generate robust, cell-level dynamic patterns remains an open question.

Here, we explore single-cell-scale dynamic patterns emerging from collective motion driven by polarized, force-balanced interactions. We focus on a minimal self-alignment framework, where the polarity of each cell aligns with its own movement, implemented within a two-dimensional (2D) vertex model of confluent tissues. In this setting, such force-balanced interactions arise from polarity-dependent interfacial tensions and simplified attractive–repulsive forces along cell–cell contacts, which drive collective motion through internal rearrangements. This minimal mechanism produces a unique, previously unreported antiparallel circulation pattern at the level of constituent cells. Guided by this prediction, we observe a similar circulation pattern in aggregating *Dictyostelium discoideum*, and a simple action–reaction argument explains how neighboring circulations adopt opposite orientations. Together, these results show that polarized force-balanced interactions provide a robust and underappreciated route for generating dynamic microscopic patterns in multicellular systems.

## II. MODEL

We now consider a confluent tissue composed of *N*_c_ cells in a 2D periodic box: 0 < *x* < *L*_*x*_ and 0 < *y* < *L*_*y*_ [Fig. 1(a)]. The tissue consists of *N*_m_ motile-type cells, indexed by ℳ ≔ {1,2, ⋯, *N*_m_} and *N*_n_ ≔ *N*_c_ − *N*_m_, and nonmotile-type cells, indexed by 𝒩 ≔ {*N*_m_ + 1, ⋯, *N*_c_}, with a mixing ratio *ϕ* ≔ *N*_m_/*N*_c_. We describe multicellular dynamics using a 2D vertex model [39– 42], specified by the vertex positions {***r***_*i*_}, and the polarity orientations of motile cells {*θ*_*α*_}_*α*∈ℳ_, where ***q***_*α*_ = (cos *θ*_*α*_, sin *θ*_*α*_).

**FIG. 1.**
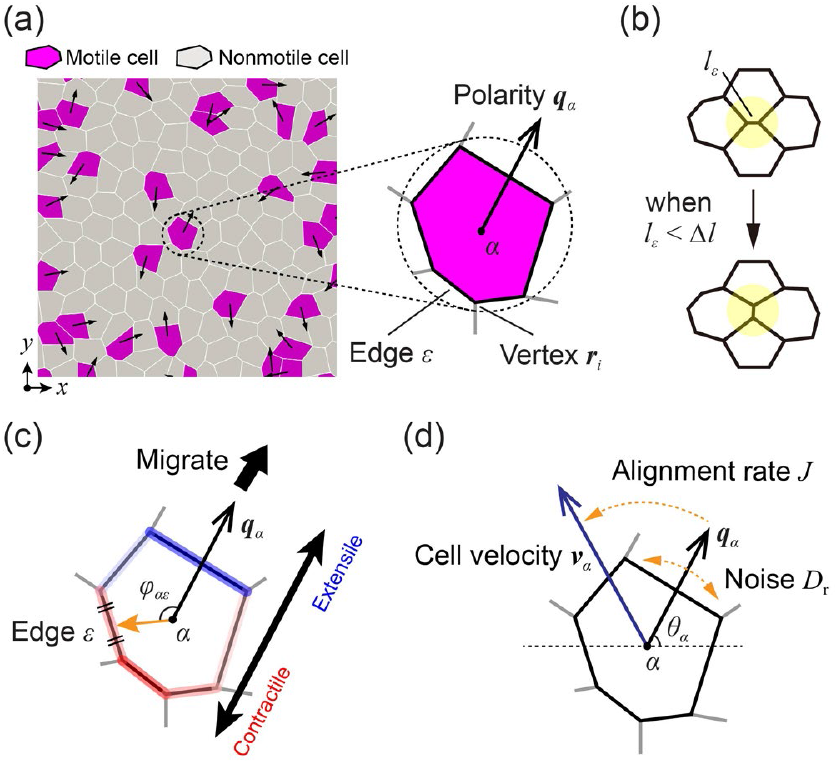
Setup of the vertex model with a self-aligned tension gradient. (a) The system is represented by a polygonal tiling with periodic boundary conditions. Each polygon expresses the shape of an individual cell. Purple and gray polygons represent motile-type and nonmotile-type cells, respectively. Arrows indicate polarity vectors of the motile-type cells. (b) Schematic illustration of a T1 transition. When the edge length, *l*_*ε*_, becomes shorter than Δ*l*, the cell configuration is rearranged by reconnecting the network of vertices and edges. (c) Tension gradient generated within a motile-type cell at its boundary with adjacent cells. Relative to the polarity vector, negative tension at the leading edge causes the edge to extend, and positive tension causes the rear edge to contract. (d) Schematic illustration of self-alignment. Each polarity vector tends to align with the velocity of the cell centroid with an alignment rate, *J*, and undergoes rotational diffusion with noise intensity, *D*_r_.

We first assume self-alignment, meaning that the polarity of each motile-type cell aligns with the direction of its own velocity, consistent with previous theoretical and experimental studies [24,26,34–37,43–46]. Additionally, accounting for rotational noise, the time evolution of the orientation of the polarity, denoted by {*θ*_*α*_}_*α*∈ℳ_, is given by

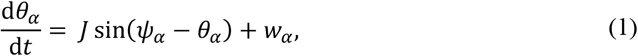

where *J* is the alignment rate and *ψ*_*α*_ is the orientation of the velocity of the *α*-th cell. Additionally, *w*_*α*_ represents white Gaussian noise, satisfying ⟨*w*_*α*_(*t*)⟩ = 0 and ⟨*w*_*α*_(*t*)*w*_*α*_′(*t*^′^)⟩ = 2*D*_r_*δ*_*αα*_′ *δ*(*t* − *t*^′^), where *D*_r_ is the noise intensity. From Eq. (1), the characteristic time scale on which polarity aligns with velocity is determined by *J* and *D*_r_.

We next describe the multicellular dynamics driven by self-aligned tension gradients. A cell monolayer is represented as a polygonal tiling, where each polygon corresponds to a cell, and vertices and edges are shared among adjacent cells [Fig. 1(a)]. Cell rearrangements occur through T1 transitions when the length of an edge becomes smaller than the threshold length Δ*l* [Fig. 1(b)].

The positions of the vertices {***r***_*i*_} evolve under overdamped dynamics,

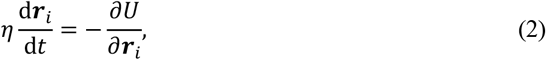

where *η* is the substrate friction coefficient. The left-hand side represents the frictional drag on vertex *i*, and the right-hand side gives the mechanical force derived from the effective energy, *U*. *U* is given by a standard vertex-model functional widely employed in previous studies [40,41,47]:

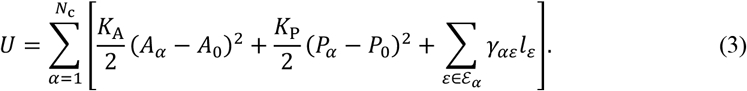

The first term represents the area elastic energy, where *K*_A_ is the area elasticity modulus, *A*_*α*_ is the area of the *α*-th cell, and *A*_0_ is the preferred cell area. The second term represents the perimeter elastic energy, with *K*_P_ the perimeter-elastic modulus, *P*_*α*_ the perimeter of the *α*-th cell, and *P*_0_ the preferred perimeter. The third term accounts for interfacial energy, where *γ*_*αε*_ is the tension on the *ε*-th edge of the *α*-th cell, *l*_*ε*_ is the length of the *ε*-th edge, and ℰ_*α*_ is the set of edges forming the *α*-th cell.

Our model assumes that each motile cell generates a driving force by establishing a polarity-aligned interfacial tension gradient along its edges, consistent with previous studies of polarity-dependent interfacial tensions in epithelial systems [29–31,48,49]. The tension, *γ*_*αε*_ for *ε* ∈ ℰ_*α*_, is defined as

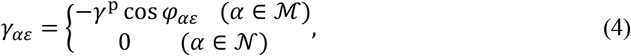

where *γ*^p^ represents the strength of polarized tension, and *φ*_*αε*_ denotes the angle between the polarity vector ***q***_*α*_ = (cos *θ*_*α*_, sin *θ*_*α*_) and the vector from the center of the *α*-th cell to the midpoint of the *ε*-th edge [Fig. 1(d)]. Through Eq. (4), each motile cell generates a gradient of interfacial tension—higher at the trailing side (cos *φ*_*αε*_ < 0) and lower at the leading side (cos *φ*_*αε*_ > 0), thereby producing a polarity-directed active force within the tissue. Although *γ*_*αε*_ depends implicitly on {***r***_*i*_} and {*θ*_*α*_} through *φ*_*αε*_, we assume that tensions are set autonomously by cellular activity rather than arising from mechanical equilibration. Thus, when calculating forces from Eq. (2), we treat *γ*_*αε*_ as constant with respect to vertex positions (i.e., *∂γ*_*αε*_/*∂****r***_*i*_ = 0). This assumption ensures that the tension gradient acts as an active force, a detailed-balance-breaking force while still satisfying force balance.

All parameters are nondimensionalized by the unit length 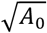, unit time *η*/(*K*_A_*A*_0_), and unit energy *K*_A_*A*_0_^2^. The parameters, *K*_p_ = 1, *γ*^p^ = 0.20, *N*_c_ = 32 × 32, and *ϕ* = 0.50, are fixed in all simulations, unless otherwise specified, while *J* and *D*_r_ are varied. To ensure that varying the values of *J* and *D*_r_ does not affect the tissue fluidity, the nondimensionalized preferred perimeter 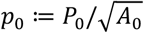, known as the target shape index, is fixed at *p*_0_ = 3.90, meaning that the system remains in a fluid-like state [50] (see Appendix A). Equations (1) and (2) are solved numerically using the Euler method from *t* = 0 to *t* = 10^4^ with a time step of Δ*t* = 0.01. The threshold length of T1 transitions is set to Δ*l* = 0.05.

## III. RESULTS

### A. Self-aligned tension gradients induce an antiparallel circulation pattern

By varying the alignment rate, *J*, and noise intensity, *D*_r_, we first found that the cells underwent phase separation when *J* > *D*_r_: motile cells spontaneously cohered and formed domains surrounded by nonmotile cells [Fig. 2(a) top row and Movie S1]. To quantify the phase separation, we introduced the separation degree, *μ*, as an order parameter,

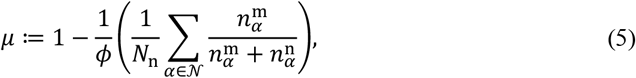

where 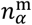 and 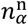 are the numbers of motile-type and nonmotile-type cells adjoining the *α*-th cell, respectively; *μ* ≈ 1 indicates phase separation, while *μ* ≈ 0 indicates a homogeneous system.

**FIG. 2.**
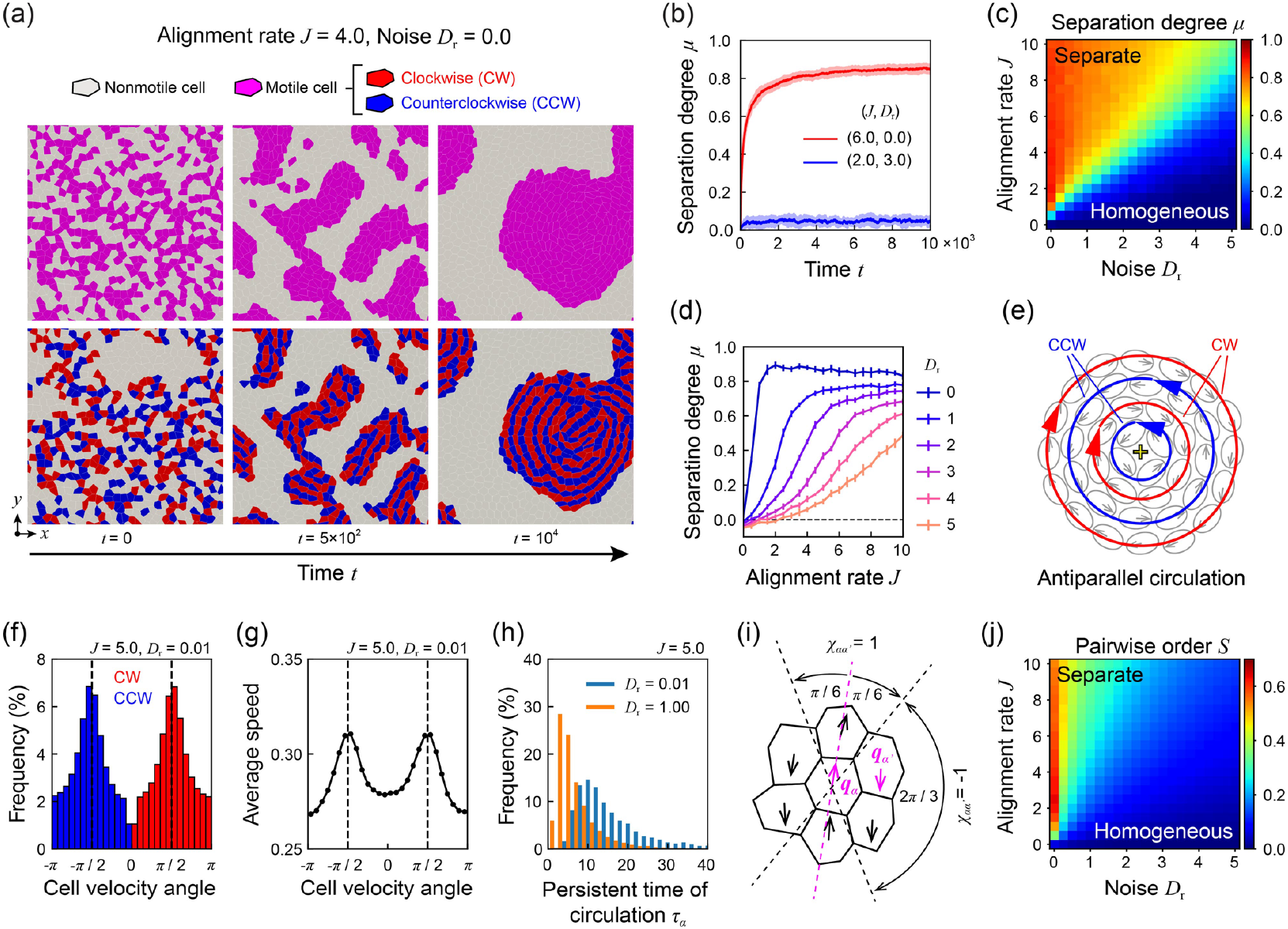
Phase separation and the emergence of antiparallel cell circulation in the vertex model with a self-aligned tension gradient. (a) Time evolution of the system with an alignment rate of *J* = 4.0 and a noise intensity of *D*_r_ = 0.0 (also shown in Movies S1 and S2). Motile-type cells rotating clockwise (CW) around the center of circulation are colored red, and those rotating counterclockwise (CCW) are colored blue. (b) Time evolution of the separation degree, *μ*, for two parameter sets: (*J, D*_r_) = (6.0,0.0) and (*J, D*_r_) = (2.0,3.0). The line and band widths in the plot indicate the statistical average values and standard deviations, respectively. (c) Phase diagram of the separation degree, *μ*, as a function of *J* and *D*_r_. (d) Separation degree, *μ*, as a function of *J* with *D*_r_ = {0.0, 1.0, 2.0, 3.0, 4.0, 5.0}. (e) Schematic illustration of antiparallel cell circulation within a motile-type cell domain. Red arrows represent CW trajectories, and blue arrows represent CCW ones. The circulations are concentrically aligned, with adjacent layers moving in antiparallel directions. (f) Distribution of cell migration directions (CW: red; CCW: blue) as a function of cell velocity angle with (*J, D*_r_) = (5.0, 0.01). The cell velocity angle is defined as the angle between the direction of cell migration and the vector from the domain center to the center of circulation. (g) Average cell speed as a function of cell velocity angle with (*J, D*_r_) = (5.0, 0.01). (h) Distribution of the circulation persistence time, *τ*_*α*_, with *J* for *D*_r_ = 0.01 (blue) and *D*_r_ = 1.00 (orange). (i) Schematic definition of the geometric factor *χ*_*αα*_′ for calculating the pairwise order parameter *S*. If the center of the *α*^′^-th cell, adjacent to the *α*-th cell, lies within the sector ranging from *π*/3 forward or *π*/3 backward relative to the polarity vector, ***q***_*α*_, of the *α*-th cell, then *χ*_*αα*_′ = 1; otherwise, *χ*_*αα*_′ = −1. (j) Phase diagram of the pairwise order, *S*, as a function of *J* and *D*_*r*_.

During phase separation, *μ* increases and eventually reaches a plateau value, whereas it remains near zero when no separation occurs [Fig. 2(b)]. The plateau value approaches unity for sufficiently large *J* and small *D*_r_ values [Fig. 2(c)].

When *J* < *D*_r_, phase separation does not occur; even when a pre-separated structure is used as the initial condition, the system reverts to a disordered configuration (see Appendix B). Separation persists even at *D*_r_ = 0 as long as *J* ≠ 0, whereas no separation is observed when *J* = 0 and *D*_r_ ≠ 0. These results demonstrate that self-alignment is essential for phase separation [Fig. 2(c)]. Moreover, for small *D*_r_, *μ* exhibits a steep transition with respect to *J*, revealing a critical point, whereas the transition becomes more gradual as *D*_r_ increases [Fig. 2(d)]. These findings indicate that phase separation arises from polarity alignment coupled to active forces and is therefore qualitatively distinct from thermally driven phase separation.

We next examined the cell movement and polarity dynamics within the domains formed under phase-separation conditions. We found that motile cells self-organize into a vortex-like pattern [Fig. 2(a) bottom row and Movie S2] (see Appendix C), characterized by concentric alignment of both velocity and polarity. Cells sharing the same velocity and polarity direction assemble into chain-like arrangements, which we refer to as circulations. These circulations are organized into concentric, onion-like layers, with adjacent circulations aligned and moving in antiparallel directions [Fig. 2(e)].

To quantify the emergence of antiparallel circulations, we performed simulations starting from a circular domain composed of motile cells with random initial polarity directions. Under a phase-separation condition with *J* = 5.0 and *D*_r_ = 0.01, we calculated the statistics of cell movement directions and speeds [Fig. 2(f) and 2(g)]. The results showed that cells predominantly move either clockwise or counterclockwise, exhibiting maximal velocity in the tangential directions, demonstrating that cells migrate in a concentric antiparallel manner. We further evaluated the persistence time of cell migration along the circulation, denoted *τ*_*α*_ (see Appendix C), for *D*_r_ = 0.01 and *D*_r_ = 1.00. This analysis revealed that cells migrate more persistently at lower noise [Fig. 2(h)], indicating that circulation persists for significant durations under low-noise conditions.

Furthermore, to quantify the emergence of antiparallel alignment between adjacent circulations, we introduce a pairwise order parameter *S*, defined as

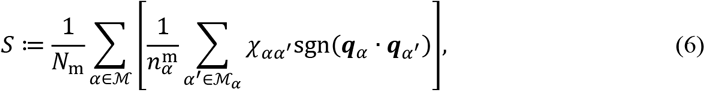

where, ℳ_*α*_ denotes the set of motile-type cells adjoining the *α*-th motile-type cell. The geometric factor *χ*_*αα*_′ is given by

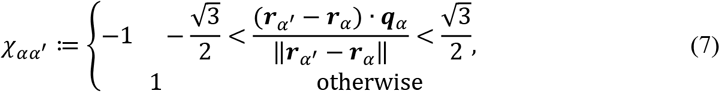

where ***r***_*α*_ is the centroid position of the *α*-th motile-type cell. Thus, *χ*_*αα*_′ = 1 when the neighboring *α*^′^-th cell lines within a forward or backward *π*/3 cone relative to the polarity direction, and *χ*_*αα*_′ = −1 otherwise [Fig. 2(i)]. Values of *S* ≈ 1 indicate predominantly antiparallel alignment within each domain, whereas *S* ≈ 0 corresponds to random polarity.

By averaging *S* over the time interval *t* = 0.95*T* to *t* = *T*, we confirm that phase separation is accompanied by the emergence of strong antiparallel alignment [Fig. 2(j)]. This alignment, and the resulting structure, emerges and remains stable under low-noise conditions, whereas it appears only transiently or disappears entirely when noise is higher. These results demonstrate that motile cells within the separated domains robustly form continuous, concentric antiparallel circulation patterns.

### B. Ordering leads to dynamic scaling with 1/3 power-law domain growth

To investigate the macroscopic behavior of the observed phase separation, we first calculate the mean-square displacement (MSD) of motile-type cells [Fig. 3(a)] (see Appendix D). At short timescales, cells exhibit ballistic motion, characterized by an MSD ∝ *τ*^2^. At longer timescales, the MSD follows a subdiffusive power law, MSD ∝ *τ*^*λ*^ with *λ* < 1, for large *J* values, indicating constrained motion within a phase-separated structure. In contrast, for small *J*, the MSD displays normal diffusion, MSD ∝ *τ*^1^, reflecting noise-dominated behavior. These results show that motile-type cells exhibit subdiffusive dynamics characteristic of phase-separating systems.

**FIG. 3.**
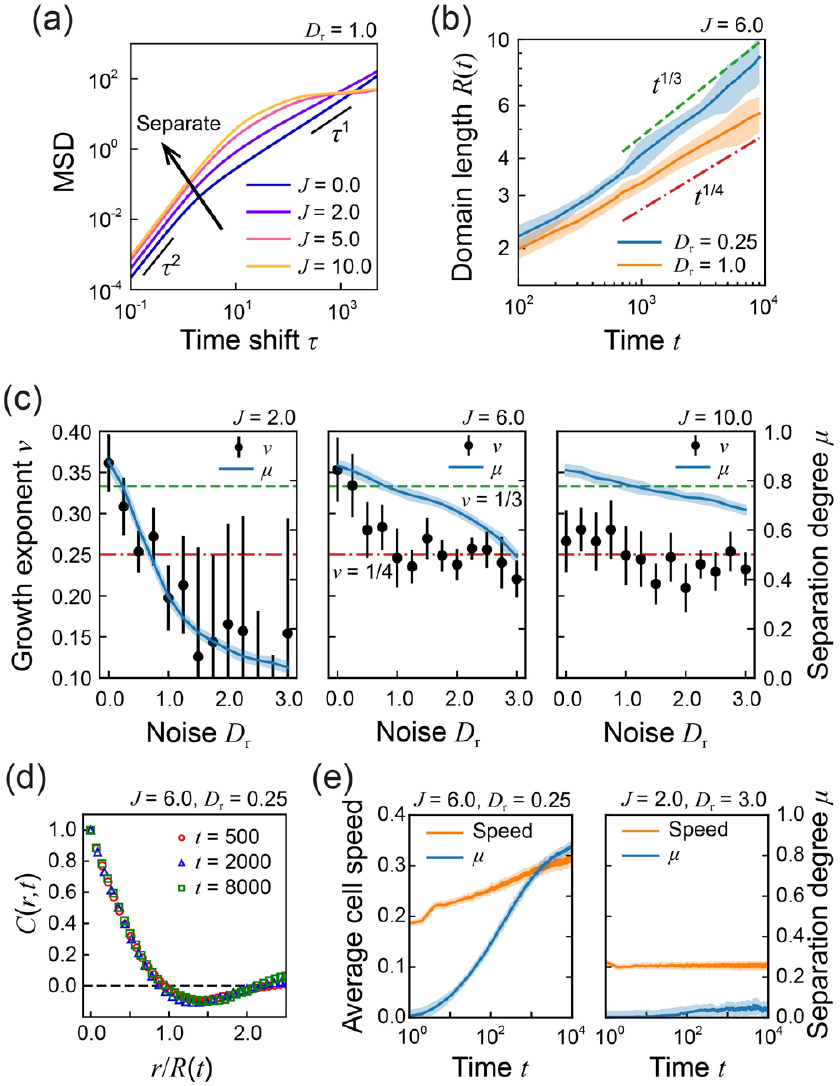
Macroscopic behavior of motile-type cells and domains. (a) Mean-square displacement (MSD) of motile-type cells as a function of the time shift, *τ*. (b) Time evolution of the domain length, *R*, with *J* = 6.0 for *D*_r_ = 0.25 (blue) and *D*_r_ = 1.0 (orange). The line and band widths in the plot indicate the statistical average values and standard deviations, respectively. (c) Growth exponent, *ν* (black point), and separation degree, *μ* (blue line), as a function of noise intensity *D*_r_ with *J* = 2.0 (left), *J* = 6.0 (middle), and *J* = 10.0 (right). The green dashed lines and red dash-dotted lines indicate *ν* = 1/3 and *ν* = 1/4, respectively. (d) Spatial correlation function of cell types, *C*(*r, t*), as a function of the distance between cells, *r*, normalized by the domain length, *R*, with (*J, D*_r_) = (6.0, 0.25). (e) Average speed of motile-type cells (orange line) and separation degree, *μ* (blue line), as a function of time, *t*, with (*J, D*_r_) = (6.0, 0.25) (left) and (*J, D*_r_) = (2.0, 3.0) (right).

We next calculated the typical domain length, *R*(*t*), to characterize the growth dynamics of the phase-separated domains [Fig 3(b)] (see Appendix D). In the parameter range where phase separation occurs, *R*(*t*) grows according to a *t*^1/3^ power law once separation is fully established, that is, when the alignment rate is moderate (*J* ≲ 7) and noise is low (*D*_*r*_ ≲ 0.5). Outside this range, the growth becomes slower, following a *t*^1/4^ power law [Fig. 3(c)]. In both the 1/3 and 1/4 regimes, spatial correlation functions at different times collapse onto a single curve when distances are rescaled by the instantaneous domain length *R*(*t*) [Fig. 3(d)]. This collapse indicates that the system obeys dynamic scaling, evolving self-similarly in time, with coarsening governed by a single characteristic growing length scale [51].

Our model exhibits a growth exponent of *ν* = 1/3 in the domain-growth law *R* ∝ *t*^*ν*^ within the low-noise regime. A similar exponent arises in motility-induced phase separation (MIPS) driven by run-and-tumble motion [52,53]. In contrast, *ν* = 1/4 is commonly observed in MIPS driven by active Brownian motion [1,53,54] and in the standard Vicsek model [55,56]. The difference in scaling exponents suggests distinct mechanisms of domain coarsening. The *ν* = 1/4 exponent is typically associated with domain coalescence mediated by diffusion–coalescence processes [57–60], although the microscopic origin of the 1/4 law in the Vicsek model remains debated. By comparison, the *ν* = 1/3 exponent is characteristic of diffusion-limited growth [61–63], indicating that in our system, domain coarsening is primarily controlled by diffusive transport along the interfaces of motile-cell domains.

Moreover, to characterize cell dynamics during phase separation, we quantified the average speed of motile cells. We found that their speed increases over time when phase separation occurs [Fig. 3(e), left], whereas it remains approximately constant in the absence of separation [Fig. 3(e), right]. This behavior indicates that the mechanism driving phase separation in our system differs fundamentally from MIPS, where aggregation arises because repeated collisions slow particles down and reduce their effective speed [53].

### C. Pairwise translational cell motion leads to persistent circulation

To clarify the elementary processes of cell circulation, we examined pairwise cell movements. We simulated the behavior of a simplified system composed of two motile-type cells, A and B, surrounded by nonmotile-type cells and found that a persistent pairwise motion of the motile-type cells emerged [Fig. 4(a) and Movie S3]. To quantify the persistence of this motion, we calculated the autocorrelation function of the average velocity of cells A and B, 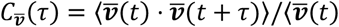. ***v***(*t*)⟩, under low-noise (*D*_r_ = 0.01) and high-noise (*D*_r_ = 1.00) conditions at *J* = 5.0. We found that the correlation time was markedly longer under low-noise conditions than under the high-noise conditions [Fig. 4(b)], consistent with the persistence-time analysis of circulation obtained earlier [Figs. 2(h) and 3(f)].

**FIG. 4.**
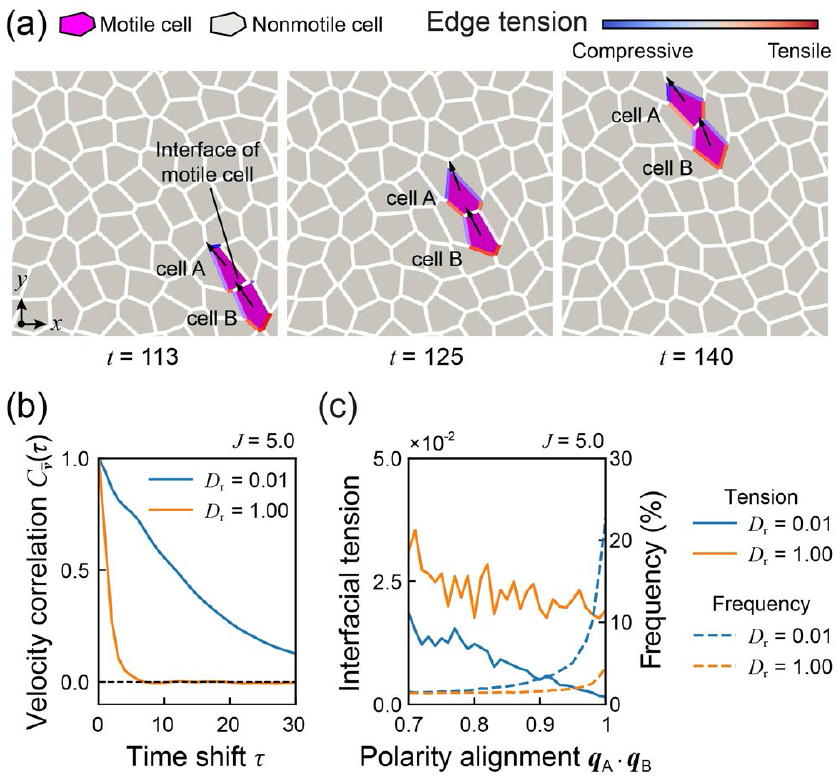
Translational motion of a pair of motile-type cells. (a) Snapshots of a motile-type cell pair in translational motion (also shown in Movie S3). Motile-type and nonmotile-type cells are colored purple and gray, respectively. Arrows indicate the polarity vectors of motile-type cells, denoted by ***q***_A_ and ***q***_B_. Edge colors indicate the intensity of edge tension. (b) Autocorrelation of the mean velocity of the motile-type cells as a function of time shift, *τ*, with *D*_r_ = 0.01 (blue) and *D*_r_ = 1.00 (orange). (c) The statistical average of the interfacial tension between the motile-type cells as a function of polarity alignment ***q***_A_ ⋅ ***q***_B_ (solid line) and the distribution of the polarity alignment (dashed line), with *D*_r_ = 0.01 (blue) and *D*_r_ = 1.00 (orange).

To understand how the paired configuration persists, we analyzed the frequency of polarity alignment between the two motile cells and their interfacial tension [Fig. 4(c)]. Alignment was assessed by the inner product of the polarity vectors of cells A and B: ***q***_*A*_ ⋅ ***q***_*B*_. Under high-noise conditions, the frequency of aligned states increases as ***q***_*A*_ ⋅ ***q***_*B*_ increases; however, even at strong alignment, the interfacial tension remains positive. In contrast, under low-noise conditions, the frequency of the aligned configuration is higher than that under high-noise conditions. Remarkably, as alignment approaches the fully aligned state (***q***_*A*_ ⋅ ***q***_*B*_∼1), the interfacial tension approaches zero, indicating that positive and negative tensions exerted by the rear and front cells cancel each other at their interface. This cancellation mechanism is consistent with a previous report of chain-like collective migration driven by a tension gradient [31].

These findings indicate that once a head-to-tail configuration forms, the pair behaves as a mechanically self-stabilizing unit. The persistence of polarity in low-noise environments prevents the cancellation of tensions from being disrupted, allowing the pair to migrate coherently until perturbed by noise, collisions, or shape fluctuations.

### D. Self-aligned tension gradient induces stable antiparallel motion of the circulation flows

To clarify how the combination of tension gradient and self-alignment gives rise to antiparallel circulation, we constructed a simple mathematical model. Specifically, to analyze the distinct contributions of tension-gradient-driven interactions from self-propelled forces, we introduced both terms into a minimal two-rail system and compared their effects.

We consider two parallel circulation rails composed of motile cells [Fig. 5(a)]. Motivated by the translational migration of motile cells observed in Fig. 4(a), we assume that within each rail, all cells share a common velocity and polarity direction. We denote the velocity and polarity of the *i*-th rail (*i* = 1,2) along the *x*-th axis by *v*_*i*_ and *q*_*i*_, respectively. The polarity *q*_*i*_ is treated as a scalar defined by *q*_*i*_ = cos *θ*_*i*_, where *θ*_*i*_ is the angle between the polarity vector and the rail direction. Each rail experiences friction against the substrate with a coefficient, *η*, fixed to the unit for simplicity. Under this setup, the motion equation of the *i*-th rail is written as

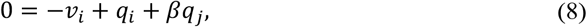

where *i, j* = 1,2 and *j* ≠ *i*. The first term represents the substrate friction force. The second term represents a polarity-dependent self-propelled force. The third term represents a reaction force from another rail, proportional to the polarity of the *j*-th rail and controlled by *β*.

**FIG. 5.**
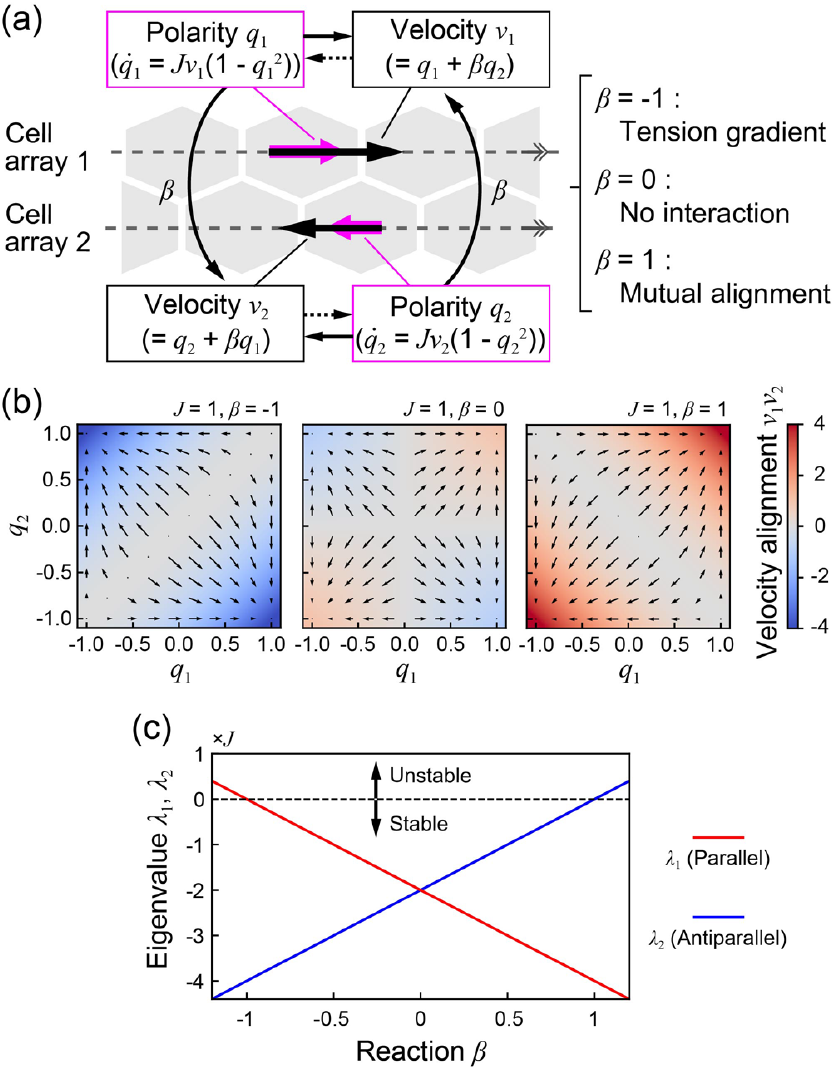
Stability of the circulation configuration in a two-rail model. (a) Setup of the two-rail model. Each dashed line represents the cell array corresponding to cell circulation. The *i*-th array has a flow velocity, *v*_*i*_ (black arrow), along the array and polarity, *q*_*i*_ (magenta arrow). The interaction between cell arrays is characterized by an interaction index, *β*. (b) The vector field of the polarities, *q*_1_ and *q*_2_, with *β* = −1 (left), *β* = 0 (middle), and *β* = 1 (right). The color indicates the value of *v*_1_*v*_2_. (c) Eigenvalues of the Jacobian matrix for fixed points corresponding to the steady parallel arrangement (red) and the steady antiparallel arrangement (blue) as a function of *β*.

To understand the basic nature of the two-rail model, we first consider the case where the velocity is aligned with polarity (sgn(*v*_*i*_) = sgn(*q*_*i*_)). In this case, Eq. (8) describes several situations that combine driving forces with polarity regulation. When *β* = 1, Eq. (8) immediately yields *v*_1_ = *v*_2_ (i.e., *q*_1_ and *q*_2_ are aligned in the same direction), corresponding to parallel motion. This situation violates force balance and corresponds to the combined case with mutual alignment and self-propelled driving forces, as defined in previous studies [17,32]. In contrast, when *β* = −1, Eq. (8) immediately yields *v*_1_ = −*v*_2_, corresponding to antiparallel motion. In this situation, the action–reaction forces generated by *q*_1_ and *q*_2_, respectively, are in force balance, and the system corresponds to the self-aligned tension-gradient case, defined in our vertex model. These results demonstrate that antiparallel circulation arises naturally from the force-balanced nature of tension gradients.

To further examine the effects of self-alignment on circulation patterns, we next incorporate polarity regulation into the simple model. The time evolution of *θ*_*i*_ is given by

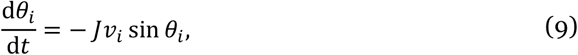

where *J* denotes the alignment rate. Combining Eqs. (8) and (9), we obtain the time evolution of the polarity *q*_*i*_:

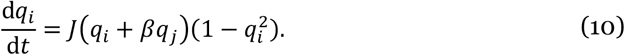

Solving Eq. (10) yields four fixed points of the polarity pair (±1, ±1) and (±1, ∓1), corresponding to the parallel and antiparallel configurations, respectively. The points that satisfy *q*_*i*_ + *βq*_*j*_ = 0 are also fixed points; however, they correspond to *v*_*i*_ = 0. Therefore, we exclude these cases from consideration.

Using Eq. (10), we determined which fixed point the dynamics converge to and evaluated the steady-state value of *v*_1_*v*_2_ in the (*q*_1_, *q*_2_) phase space [color-coded in Fig. 5(b)]. Moreover, to investigate the linear stability of the fixed points, we analytically calculated the eigenvalues *λ*_1_ and *λ*_2_ of the Jacobian matrix **J** of Eq. (10) evaluated at each fixed point. We found that *λ*_1_ and *λ*_2_ reflect the stabilities of the parallel and antiparallel configurations, respectively. The Jacobian matrix is **J** = *J*(−2*β* − 2)**I** at (*q*_1_, *q*_2_) = (±1, ±1) and **J** = *J*(2*β* − 2)**I** at (*q*_1_, *q*_2_) = (±1, ∓1), where **I** denotes the identity matrix. Thus, the eigenvalues depend on *β* as *λ*_1_ = *J*(−2*β* − 2) and *λ*_2_ = *J*(2*β* − 2) [Fig. 5(c)].

When the driving force is purely tension-gradient-based (*β* = −1), the polarity dynamics converge to the antiparallel configuration [Fig. 5(b), left]. In this case, the antiparallel configuration is linearly stable (*λ*_2_ = −4*J* < 0), whereas the parallel configuration is not (*λ*_1_ = 0) [Fig. 5(c)]. This behavior is consistent with our vertex model, in which force-balanced tension gradients combined with self-alignment give rise to antiparallel circulation. In contrast, when motion is driven solely by self-propulsion (*β* = 0), the dynamics can converge to either the parallel or antiparallel configuration depending on initial conditions [Fig. 5(b), middle], and both are equally stable (*λ*_1_ = *λ*_2_ = −2*J* < 0) [Fig. 5(c)]. Finally, when the interaction corresponds to mutual alignment (*β* = 1), the polarity dynamics converge to the parallel configuration [Fig. 5(b), right]; the parallel configuration is linearly stable (*λ*_1_ = −4*J* < 0), whereas the antiparallel configuration is not (*λ*_2_ = 0) [Fig. 5(c)]. Therefore, antiparallel patterns do not emerge through mutual alignment alone.

These results demonstrate that the antiparallel pattern is a characteristic feature emerging from self-aligned, force-balanced driving forces, distinct from dynamics governed by self-propulsion or mutual alignment. Importantly, this mechanism is not limited to the vertex-model framework or to a specific tension-gradient-driven interaction; rather, it may apply more generally when constituents form head-to-tail units and interact through locally force-balanced couplings under self-alignment.

### E. *D. discoideum* demonstrates an antiparallel circulation pattern

To examine whether antiparallel circulation arises in biological systems, we analyzed the collective cell movements of *D. discoideum*. These cells are monopolar and motile, and they exhibit vortex-like motion [20] even in the absence of chemotactic signals [64]. Their polarity is established through front-to-back cell–cell contacts [64], which is consistent with the polarity–tension coupling assumed in our model. This suggested that an antiparallel circulating state might exist but remain overlooked in three-dimensional environments, where cell trajectories are more complex. To clearly identify cell movements, we confined differentiated cells to quasi-2D geometry by seeding cells in a custom-made 2.5-μm-thick chamber placed between a glass coverslip and polydimethylsiloxane [Fig. 6(a)] (see Appendix E) [65,66]. The cells formed a disk-like aggregate [Fig. 6(b)], within which we observed a circulation pattern strikingly similar to that predicted by our simulations [Fig. 6(c) and Movie S4].

**FIG. 6.**
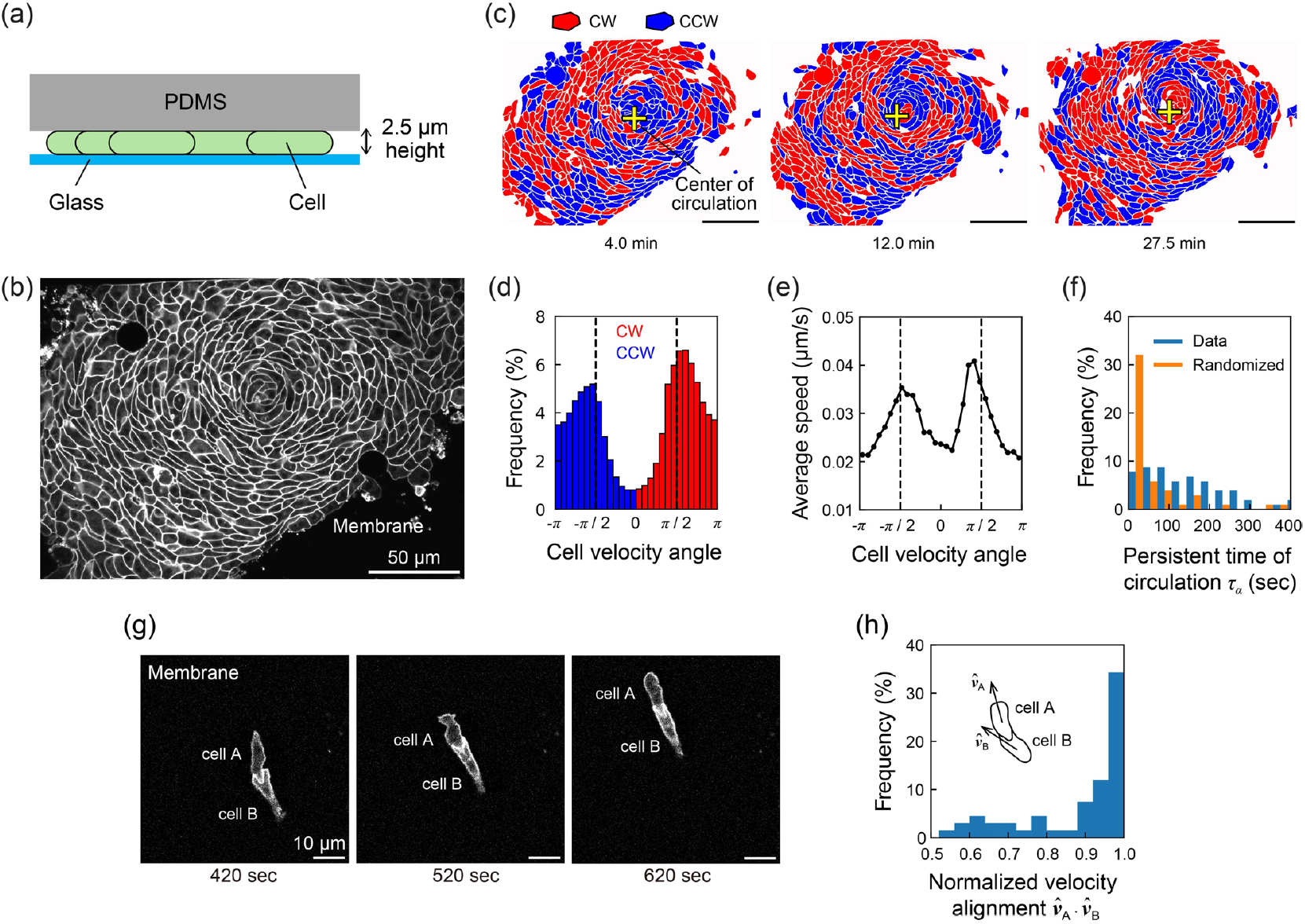
Antiparallel circulation and translational motion of *Dictyostelium discoideum* cells. (a) Experimental setup for the observation of *D. discoideum*. (b) A snapshot of a *D. discoideum* aggregate in a 2D chamber. (c) Time evolution of the aggregate with circulation directions of cells around the center (also shown in Movie S4). Cells exhibiting clockwise (CW) circulation are colored red, and those with counterclockwise (CCW) circulation are colored blue. The yellow crosses mark the center of circulation. (d) Distribution of cell migration directions (CW: red; CCW: blue) as a function of cell velocity angle. (e) Average cell speed as a function of cell velocity angle. (f) Distribution of the circulation persistence time, *τ*_*α*_, for the raw data (blue) and randomized data generated via bootstrap sampling (orange). (g) Snapshots of a pair of *D. discoideum* cells in translational motion (also shown in Movie S5). (h) Distribution of the normalized velocity alignment, ***, of a pair of *D. discoideum* cells.

To quantify this behavior, we measured the direction, speed, and persistence time of cell motion using the same definitions applied in our simulations [compare Fig. 6(d)–(f) with Fig. 3(f)–(h)]. For persistence time, we further evaluated the significance by comparing the observed data with randomized trajectories generated by bootstrap resampling. These analyses show that the quantitative features of cell motion in the aggregates are qualitatively consistent with those from our model. This finding demonstrates that *D. discoideum* cells exhibit continuous, concentric antiparallel circulation patterns similar to those predicted in our simulations. We note that the circulations were not always single-cell-wide lanes but sometimes consisted of several cells, likely reflecting side-by-side adhesion [67], an effect not included in our model, leading to broader circulation bands than those observed in the vertex simulations.

Furthermore, when *D. discoideum* cells were seeded at low density, we frequently observed persistent translational motion in which two cells aligned in a head-to-tail configuration and migrated together as a pair [Fig. 6(g) and Movie S5], comparable to the configuration observed in our simulations [Fig. 4(a)]. We quantified this translational alignment using the inner product of the normalized velocity vectors of cells A and B, 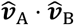 [Fig. 6(h)]. Fully aligned states 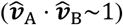 were frequent, consistent with the simulations [Fig. 4(c)]. This configuration persists until the pair encounters another cluster—typically a large one, at which point the pair merges or reorganizes. This behavior resembles the migration patterns of 2D cell clusters under an imposed gradient of the chemoattractant, cAMP [64]. These results demonstrate that *D. discoideum* cells can induce a head-to-tail configuration as a mechanically self-stabilizing unit, suggesting that a similar mechanism predicted by the model could underlie the antiparallel circulations observed in *D. discoideum*.

## IV. DISCUSSION

Our results show that antiparallel cell circulation can emerge from polarized, force-balanced interactions, providing a distinct alternative to the widely studied picture of self-propelled cells driven by substrate traction. In many active-matter models of multiple cells, motility is represented as a polarity-dependent self-propulsion force that is balanced by friction at the cell–substrate interface. In our framework, in contrast, the driving forces are polarity-dependent interfacial forces (or, more generally, polarized cell–cell interactions) that act along contacts between cells. These forces are locally force-balanced, where each interface exerts equal and opposite forces on its neighboring cells—yet their polarization, together with polarity–velocity alignment, generates sustained microscopic circulation. This mechanism shows that, when combined with self-alignment, alternative driving modes with different patterns of force transfer between neighboring cells can generate robust dynamic patterns at the cell level in multicellular systems.

This difference in how forces are applied is reflected in the emergent patterns. Existing self-alignment and mutual alignment, including Vicsek-type models, can generate vortices [68–70], unidirectional circulation [71,72], or flocking states [16–19], but the circulation within each structure is typically single-directional and often relies on center-of-mass propulsion or geometric confinement. In contrast, our model produces interdigitated antiparallel circulation, where neighboring lanes of cells move in opposite directions while the tissue remains confluent and mechanically compact. To the best of our knowledge, such constituent-level antiparallel circulation has not been reported in models based on MIPS, mutual alignment, or conventional self-alignment rules. This highlights that polarized force-balanced interactions, combined with self-alignment of polarity to velocity, define a qualitatively different route to organized flow patterns in cell collectives.

The same mechanism that produces antiparallel circulation also leads to a distinct form of phase separation. In MIPS, clusters form because particles repeatedly slow down upon encountering dense regions; reduced speed promotes accumulation. In our system, the opposite trend appears as the separation degree, *μ*, increases [Fig. 2(b)], the speed of motile cells also increases [Fig. 3(e)]. Phase separation is therefore circulation-driven, rather than slowdown-driven, establishing positive feedback between microscopic circulation and mesoscopic domain growth. Moreover, although mutual alignment often exhibits phase separation through large density fluctuations reminiscent of gas–liquid separation [73], our system maintains nearly constant density within compact domains, and domain growth is constrained by diffusion [Fig. 3(b) and 3(c)]. We refer to this distinct mechanism as circulation-induced phase separation (CirPS).

An important aspect of our results is that the mechanism is robust to how polarized force-balanced interactions are realized. We first introduced them through polarity-biased interfacial tensions but then showed that simplified force-balanced attractive–repulsive interactions between neighboring cell rails can generate qualitatively similar antiparallel circulations. This indicates that the phenomenon does not depend on the detailed form of interfacial tension; instead, the combination of (i) polarity bias and (ii) local force balance at cell–cell contacts is essential. We therefore expect that other realizations of polarized force-balanced interactions, for example, those arising from active stress patterns or directed shape changes, may likewise produce circulation-driven pattern formation.

The cellular behaviors underpinning CirPS—persistent flows and local rearrangements—are frequently observed in multicellular systems, suggesting that similar mechanisms may operate *in vivo*. In our experiments, we observed antiparallel circulations in 2D aggregates of *D. discoideum* [Fig. 6], consistent with the model prediction. Cell reorientations are thought to be important for breaking up *Dictyostelium* aggregates and regulating aggregate size and sorting; our findings raise the possibility that antiparallel circulation and CirPS contribute to these processes. Similar circulation-based mechanisms could also influence the breakup or remodeling of tumor clusters and other dense cell assemblies. Overall, our work identifies polarized force-balanced interactions as a minimal and robust route to antiparallel cell circulation and CirPS, broadening the landscape of active-matter mechanisms that can underlie organization and remodeling in multicellular systems.

## DATA AVAILABILITY

The data that support the findings of this article are openly available [citation number to be inserted here].

## ACKNOWLEDGEMENTS

We thank all the members of the Okuda Laboratory for their discussions. This work was supported by the WISE Program for Nano-Precision Medicine, Science, and Technology of Kanazawa University, Ministry of Education, Culture, Sports, Science and Technology (MEXT); the Japan Science and Technology Agency (JST), CREST [Grant No. JPMJCR1921, JPMJCR24B2, JPMJCR1923]; the Japan Society for the Promotion of Science (JSPS), KAKENHI [Grant No. 21K15081, 22H05170, 23H00384, 23H04304, 24H01398, 24H01937, 25K01118, 25H01363, 25H01771, 25K22468]; the Japanese Agency for Medical Research and Development (AMED), a Program for Technological Innovation of Regenerative Medicine Grant [23bm0704065h0003], and the World Premier International Research Center Initiative, MEXT, Japan.

## Appendices

## APPENDIX A EFFECTS OF TISSUE FLUIDITY ON PHASE SEPARATION

Varying cell properties, such as deformability, motility, and noise, can induce a jamming transition, during which tissue fluidity changes drastically [50,74–76]. To investigate the effects of tissue fluidity on phase separation, we first reproduced the jamming transition using our model. Here, we considered a tissue composed entirely of motile-type cells without self-alignment. In this model, the shape index *p*_0_, the strength of the tension gradient *γ*^p^, and the noise intensity *D*_r_ can be used to tune tissue fluidity, which corresponds to cell deformability, motility, and noise. For simplicity, the noise intensity was fixed at *D*_r_ = 1, and the mixing ratio and alignment rate were set to *ϕ* = 1 and *J* = 0, respectively. To quantitatively assess tissue fluidity, we calculated the effective diffusion coefficient *D*_eff_ ≔ *D*_s_/*D*_0_, where *D*_s_ is the self-diffusion coefficient defined as *D*_s_ ≔ MSD(*τ*)/(4*τ*) averaged for *τ* > 0.45*T*, and *D*_0_ is the free diffusion coefficient given by *D*_0_ ≔ 2*π*^2^(*γ*^p^/*η*)^2^/*D*_r_ [31,50] (see the Supplemental material [77] for the detailed calculation of *D*_0_). Here, MSD(*τ*) denotes the mean-square displacement of motile-type cells at a time shift of *τ* (see Appendix C). We classified systems with *D*_eff_ < 10^−3^ as being a solid-like state and those with *D*_eff_ ≥ 10^−3^ as fluid-like. As a result, we found that tissue fluidity transitioned between solid-like and fluid-like states depended on the values of *p*_0_ and *γ*^p^, indicating the occurrence of a jamming transition [Fig. 7(a)].

**FIG. 7.**
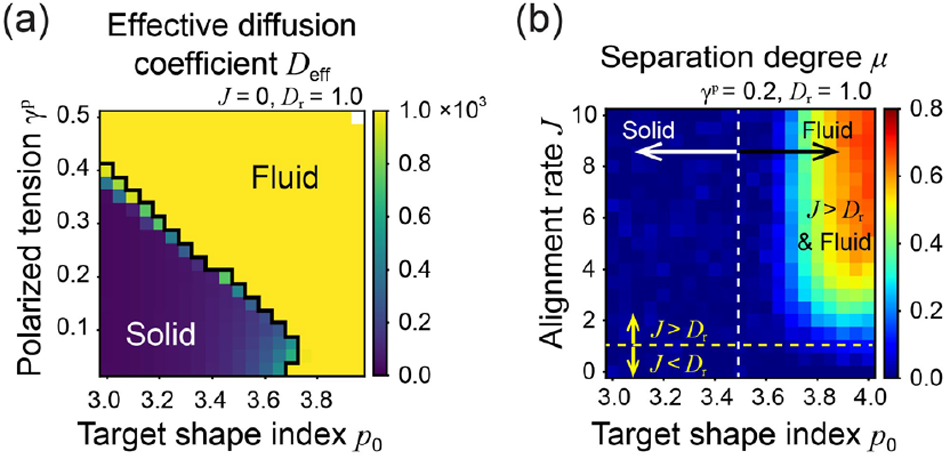
Jamming transition of the system. (a) Phase diagram of the effective diffusion coefficient, *D*_eff_, as a function of the target shape index, *p*_0_, and strength of polarized tension, *γ*^p^, with (*J, D*_r_) = (0.0, 1.0). (b) Phase diagram of the separation degree, *μ*, as a function of *p*_0_ and *J* with *γ*^p^ = 0.2 and *D*_r_ = 1.0. The white dotted line indicates the target shape index threshold, which determines whether the tissue is in a solid-like state or fluid-like state. The yellow dotted line indicates the alignment rate at which the value is equal to the noise.

We next considered a mixture of motile-type cells with self-alignment and nonmotile-type cells. In this case, the mixing ratio and noise intensity were set to *ϕ* = 0.5 and *D*_r_ = 1. By varying *p*_0_ and *J*, we observed that phase separation occurred only when the tissue was in a fluid-like state [Fig. 7(b)]. Based on these findings, all numerical simulations presented in the main text were performed under fluid-like conditions, as phase separation does not occur in the solid-like state.

## APPENDIX B DEPENDENCE ON THE INITIAL CONDITIONS

To investigate the dependence of phase separation on the initial conditions, we conducted simulations using both homogeneous and pre-separated initial states. The separated state was prepared by placing a circular domain of motile-type cells at the center of the system. When *J* > *D*_r_, phase separation occurred regardless of the initial condition, although the degree of separation, *μ* varied depending on the initial state [Fig. 8(a)]. In contrast, when *J* < *D*_r_, the system eventually relaxed into a homogeneous state, irrespective of the initial condition, and *μ* converged to zero [Fig. 8(b)]. These results indicate that the occurrence of phase separation is not influenced by the initial condition. However, when phase separation does occur, the degree of separation is influenced by the initial condition.

**FIG. 8.**
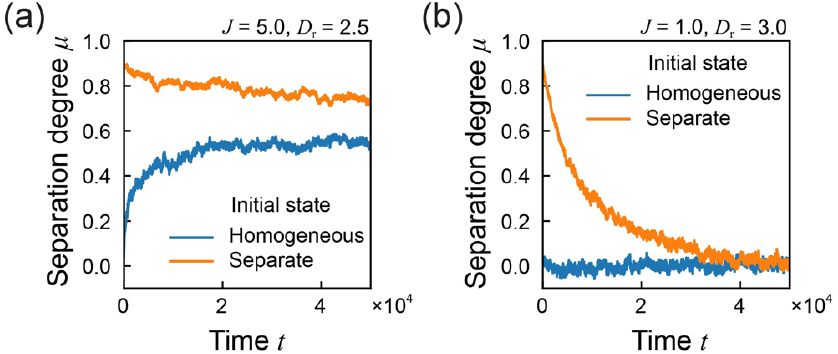
Dependence of phase separation on initial conditions. (a, b) Time evolution of the separation degree, *μ*, with (*J, D*_r_) = (5.0, 2.5) (a) and (*J, D*_r_) = (1.0, 3.0) (b). In panels (a) and (b), the blue and orange lines indicate the case with a homogeneous initial state and the case with a separated initial state, respectively.

## APPENDIX C CALCULATION OF THE CIRCULATION VELOCITY AND THE PERSISTENT TIME OF THE CIRCULATION DIRECTION

To quantify the antiparallel circulation pattern, we calculated the speed and direction of motile-type cell motion with respect to the center of circulation 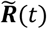 for each domain, which was determined from the velocity profile at each time *t* as follows. Using the velocity vector, ***v***_*α*_ and the position vector, 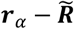, which connects the circulation center and the centroid of the *α*-th cell, 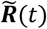, is defined as the point ***R***′ that minimizes the following function,

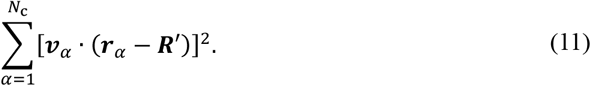

The optimal circulation center, 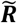 can be written explicitly as

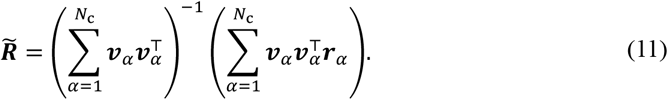

Next, the circulation velocity 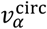 of the *α*-th cell at each time point was calculated as the projection of its velocity vector onto the tangential direction of the circular path. This is defined as

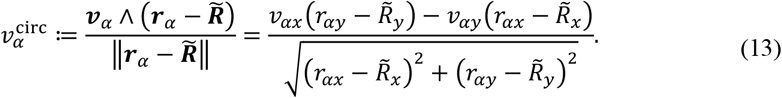

Here, 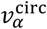 is positive when the cell is circulating in the clockwise direction. Figures 2(a) and 4(c) show the spatial pattern of the circulation direction, as indicated by the sign of 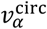 at a single time point. These findings clearly exhibit an antiparallel circulation pattern.

To investigate the frequency of the reversal movement of the cells, we calculated the persistent time of the circulation direction. Using the autocorrelation function

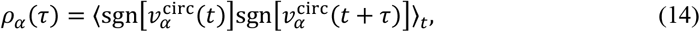

the persistent time, *τ*_*α*_ is defined as the first zero-crossing point, i.e.,

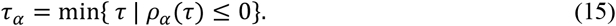

## APPENDIX D MACROSCOPIC BEHAVIOR OF THE CELLS AND DOMAINS

We calculated the MSD of motile-type cells during the time interval, *T*/2 ≤ *t* ≤ *T* as follows,

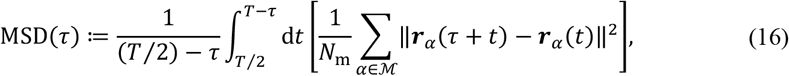

where *τ* is a time shift within the range, *τ* ∈ [0, *T*/2). For short timescales, the ballistic diffusion regime (MSD ∝ *τ*^2^) expanded as the system transitioned from a homogeneous to a phase separation state by varying the alignment rate [Fig. 3(a)]. This expansion is attributed to the increased persistence of motile-type cells when *J*/*D*_r_ > 1 [44].

For longer timescales, we observed subdiffusive behavior, characterized by MSD ∝ *τ*^*λ*^ with *λ* < 1, when *J* was sufficiently large, indicating restricted motion of motile-type cells under complete phase separation. In contrast, when *J* was small, we found normal diffusion behavior, characterized by MSD ∝ *τ*^1^, where motion was primarily governed by noise.

Next, we examined the time evolution of the typical domain length. To simplify the calculation, we first define the “existence probability” *f*_*α*_(***x***, *t*) of the *α*-th cell at coordinate ***x*** as

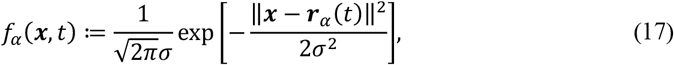

where *σ* is the standard deviation of *f*_*α*_, regarded as a spatial scale of a single cell. We set 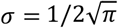 for all cells, which is equal to half of the radius of the circle with unit area. Using *f*_*α*_, we introduced the distribution function of motile-type cells, *f*(***x***, *t*), defined as

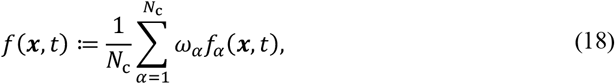

where *ω*_*α*_ is given by

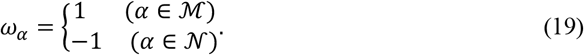

Note that *f* can be negative when nonmotile-type cells are near coordinate ***x***. Using *f*, we defined the spatial correlation function of cell types *C*(*r, t*) [63,78,79] as

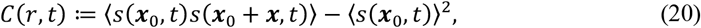

where *r* = ‖***x***‖ and *s*(***x***, *t*) = sgn *f*(***x***, *t*). The average ⟨·⟩ is taken over the coordinate ***x***_0_ and the displacement ***x*** with the given distance of *r* = ‖***x***‖. The domain length *R*(*t*) is defined as the first zero-crossing point of the correlation function, *C*(*r, t*), i.e.,

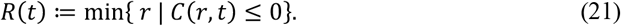

We first confirmed that *C*(*r, t*) for different time points converged to the same curve as a function of *r*/*R*(*t*) [Fig. 3(d)]. Furthermore, domain length follows *R* ∝ *t*^*ν*^ with growth exponent *ν*, which indicates the mechanisms of domain growth. Under conditions in which the noise intensity is very low (*D*_r_ ≲ 0.25) and the alignment rate is moderate (*J* ≲ 7), *ν* ≈ 1/3 [Figs. 3(b) and 3(c)], suggesting that the domain growth is governed by diffusion [62]. As the noise intensity or the alignment rate increases (*J* ≳ 7), the growth exponent rapidly shifts to *ν* ≈ 1/4 [Figs. 3(b) and 3(c)], suggesting that the domain growth is dominated by direct domain coalescence (diffusion-coalescence) processes [57–60].

**APPENDIX E: OBSERVATION OF *DICTYOSTELIUM DISCOIDEUM* AGGREGATION**

The *D. discoideum* Ax4 cell line carrying the plasma membrane marker PKBR1(N150)-mScarlet-I-2x at the *act5* locus [66] was used. The cell culture, development, microfabrication, and observation were performed as described previously [64,66]. To image a monolayer cell population [Fig. 6(b)], cells dissociated from the slugs were suspended at a density of 2.2 × 10^5^ cells/mL in phosphate buffer (PB: 12 mM KH_2_PO_4_, 8 mM Na_2_HPO_4_, pH 6.5). To observe the translational motion of two cells [Fig. 6(g)], cells were suspended at a density of 5 × 10^4^ cells/mL in PB. Then, 0.4 mL of cell suspension was loaded into a microfluidic chamber with a height of 2.5 µm [Fig. 6(b)] or 4 [Fig. 6(g)]. Fluorescence images were acquired at 22°C using an inverted microscope (IX83, Olympus/Evident) equipped with a multibeam confocal scanning unit (CSU-W1, Yokogawa) and CMOS cameras (ORCA-FusionBT, Hamamatsu Photonics).

The membrane images of *D. discoideum* were collected at 5-second intervals to obtain *I* = 361 frames [Fig. 6(b)]. The raw images were binarized using Trainable Weka Segmentation in Fiji, and cells were detected by connected component labeling. Hereafter, cells were represented by their centroid positions. A pair of cells in consecutive frames was identified as the same cell if the opponent had the smallest distance among other cells in the same frame. In this way, we obtained *N*_c_ = 507 trajectories 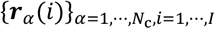, which included more than 100 frames. Note that some cells were occasionally not detected so that a true trajectory with full length could be divided into multiple trajectories with shorter lengths. Then, cells were assigned with velocity vectors, {***v***_*α*_(*i*)} at each time point. We applied a mean filter with a length of 10 frames before taking the difference between consecutive centroid positions to remove the noise. For the antiparallel circulation pattern analysis, the circulation velocity and persistent time of the circulation direction were calculated at each time point *i* (see Appendix C).

